# The impact of HIV-1 within-host evolution on transmission dynamics

**DOI:** 10.1101/233130

**Authors:** Kristof Theys, Pieter Libin, Andrea-Clemencia Pineda-Pena, Ann Nowe, Anne-Mieke Vandamme, Ana B Abecasis

## Abstract

The adaptive potential of HIV-1 is a vital mechanism to evade host immune responses and antiviral treatment. However, high evolutionary rates during persistent infection can impair transmission efficiency and alter disease progression in the new host, resulting in a delicate trade-off between within-host virulence and between-host infectiousness. This trade-off is visible in the disparity in evolutionary rates at within-host and between-host levels, and preferential transmission of ancestral donor viruses. Understanding the impact of within-host evolution for epidemiological studies is essential for the design of preventive and therapeutic measures. Herein, we review recent theoretical and experimental work that generated new insights into the complex link between within-host evolution and between-host fitness, revealing temporal and selective processes underlying the structure and dynamics of HIV-1 transmission.

## 1 Introduction

Despite the remarkable progress of the last three decades, Human Immunodeficiency Virus type 1 (HIV-1) remains a global health threat with almost two million new infections and one million AIDS-related deaths in 2016 (UNAIDS Factsheet 2017). HIV-1, as many RNA viruses, is characterized by high rates of evolutionary changes during the course of infection [1, 2, 3], and this potential for adaptive evolution has proven to be a cornerstone mechanism of viral escape from host immune responses and antiretroviral treatment (ART) [4, 5]. Yet, drug resistance rates are stabilizing or decreasing due to more efficacious and tolerable regimens, particularly in developed countries, resulting in a nearly normal life expectancy for treated HIV-1 infected individuals [6, 7].

By contrast, as immune-driven viral control by means of a preventive or therapeutic vaccine remains challenging [8], main priorities with respect to public health now concern the formulation and implementation of efficient and effective prevention strategies. The widespread implementation of Treatment as Prevention (TasP) is considered one of the most important intervention strategies to reduce the rate of HIV-1 transmission [9]. Furthermore, methodological improvements to infer transmission networks and to model epidemics in silico enable authorities to monitor dynamics of epidemic spread and ameliorate prevention strategies [10]. Despite these advances, the resulting worldwide decline in HIV-1 infections is considered insufficient to meet the UNAIDS visionary goal to end AIDS by 2030 [6]. Periodical reports of increasing incidence (e.g. Eastern Europe and Central Asia) and persistent challenges (e.g. late presenters, suboptimal adherence and increasing rates of drug resistance in Africa) illustrate the continuous need to optimize targeted intervention strategies [11, 12, 13, 6].

To this end, novel perspectives on the impact of HIV-1 within-host evolutionary processes can be of high value to better understand and mitigate population level transmission. Knowledge on the association between viral genotype and epidemic potential is crucial to formulate prevention strategies directed towards sub-epidemics that have the largest impact on the incidence rate. Furthermore, insights into the different mechanisms and virus diversity that dominate transmission and early infection will improve transmission history inference as well as vaccine design.

HIV-1 is characterized by extensive genetic diversity within and between hosts, but distinct processes and rates shape viral evolution at these levels [2, 3, 14, 15]. While within-host evolutionary dynamics are dominated by selective forces and competitive fitness (Figure 2) [2], between-host evolution is considered to be largely shaped by neutral processes and multiple HIV-1 variants co-exist at the epidemiological level, although recent findings might imply more selective action at this level than generally assumed [16, 17]. Circulating viral diversity at population level is an intrinsic reflection of transmission dynamics within the epidemic [2], but virus and host genetics additionally impact between-host dynamics [10, 18, 19, 5, 20, 2, 21]. As a result, a complex interplay of multi-scale evolutionary processes exists and the selective advantage of viral traits differs at the within-host and between-host level as conflicting evolutionary forces apply [22, 23]. While viral spread in a population is expected to select for traits that maximize transmission efficiency, viral strategies that favour within-host fitness do not necessarily benefit epidemiological fitness, leading to a delicate evolutionary trade-off between virulence and transmission [24, 25].

This review discusses the connection between HIV-1 within-host evolution and transmission dynamics and highlights recent theoretical and experimental work bridging different scales (Figure 1). We first address the processes that shape HIV viral evolution within the host, then we focus on the bottleneck transmission event and finally on how this can be translated at the population level. We end by shedding light on how these factors can affect the epidemiological transmission studies. Understanding the link between within-host processes, transmission fitness and epidemic spread will be essential to achieve HIV-1 eradication.

**Figure 1:**
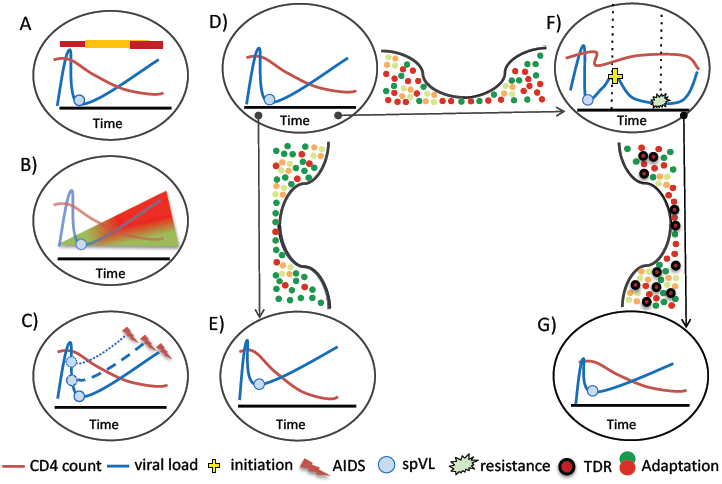
Schematic representation of a connection of within-host evolution with HIV-1 transmission. A) HIV-1 infectiousness (upper bar) maximizes with increasing viral load. B) Over the course of infection, HIV-1 increasingly (red) adapts to its environment with transient release of ancestral viruses from latent reservoirs (green). C) spVL varies between individuals, and higher spVL is associated with faster progression to AIDS. D)-G) Example of a transmission network with three transmission bottleneck events. E) Transmission occurred early in infection of ancestral viruses with limited adaptation (green) to the donor host. F) Transmission occurred late infection of variants adapted (red) to the donor host. ART initiation resulted in a decrease of viral load until the emergence of drug resistance. G) Transmission at the time of ART failure results in the transmission of drug resistance (TDR), which could affect viral load in the recipient host.

**Figure 2:**
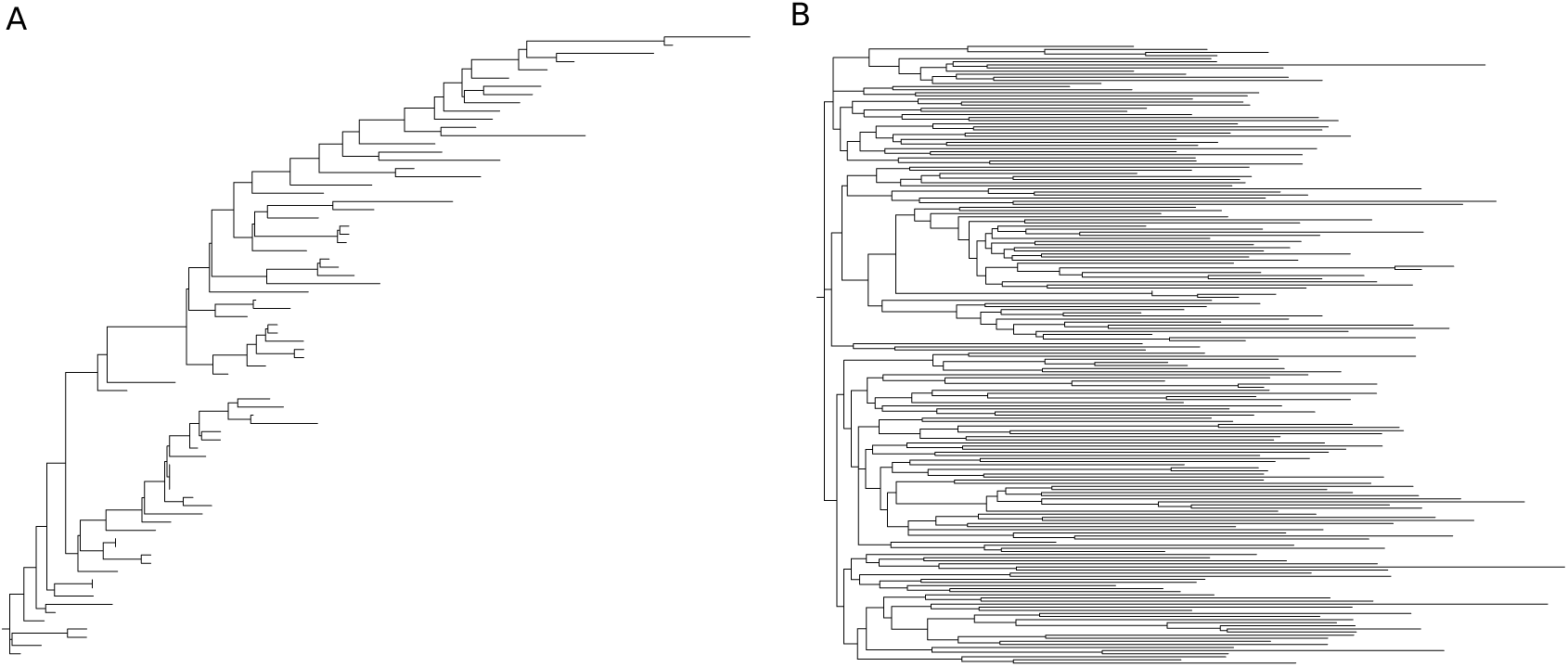
Panel (A) shows a tree consisting of HIV-1 sequences that belong to a single patient, as presented in [1]. Panel (B) shows a tree inferred from a random selection of Portuguese HIV-1 subtype B sequences, a subset of the tree presented in [90]. The within-host tree clearly demonstrates a ladder-like topology, while the population tree reflects a star-like topology.

## 2 Within-host evolutionary dynamics

Evidence of viral traits that directly promote HIV-1 transmission from one host to another is limited [26, 27]. Nonetheless, viral strains differ in their transmissibility, albeit mostly as a consequence of their effect on the different stages of disease progression.

The natural course of HIV-1 infection generally starts with an acute, primary phase followed by a chronic, asymptomatic phase which progresses into AIDS and eventually death [1]. The number of viral RNA copies in peripheral blood (viral load) and CD4+ T-cell lymphocytes (CD4 count)are routinely used to monitor the rate of disease progression. Virus load peaks soon after transmission and then levels off to a set-point viral load (spVL). spVL announces the start of the chronic phase and remains relatively stable for years [28]. AIDS eventually results from immune deterioration due to chronic immune activation and gradual depletion of the CD4 count, but rates of disease progression vary significantly among individuals [1, 2, 3, 29].

The transmission probability per contact is closely related to the viral load. HIV-1 infectiousness is therefore not constant over the time of infection, with the acute phase as an important time window [30]. A large variability in spVL has been observed between individuals, which is additionally important as spVL positively correlates with both the rate of disease progression and the per-contact transmission rate [31, 25, 32]. Recent findings suggest that within-host evolution only has a small impact on spVL variation. By contrast, changes in virus replication capacity and adaptive evolution have been linked to viral loads in early and late infection as well as to CD4 count decline [33, 34]. Others have reported that, early during infection, high levels of replication are an important marker of disease progression independently of the viral load [35, 36,5], adding to the complexity of the processes underlying HIV-1 pathogenicity and transmissibility. Coinciding with increasing viral loads, virus replication during the chronic phase continuously drives a diversifying and diverging virus population [1, 37].

While mutations are randomly generated, HIV-1 sequence space is limited by functional and immunological constraints [14], and evolutionary dynamics are governed by competitive fitness [38, 1, 39, 40]. The emergence of escape mutations illustrates adaptation to selective pressures from host immune responses and ART [5, 41]. Despite their beneficial effect in a given environment, escape mutations can confer a replication cost, with implications for viral load and transmissibility. Yet, viral fitness can be restored by epistatic interactions and compensatory mutations [42, 43, 44].

The evolutionary rate of HIV-1 can also be elevated by recombination, which is an essential mechanism to mitigate deleterious mutational load [45]. Recombination encompasses template switching between RNA strands during reverse transcription, and can lead to a new recombinant form in case of dual infection with distinct subtypes. The increasing worldwide prevalence of circulating recombinant forms could imply a replication or transmission advantage of some of these strains [46, 15].

Evolutionary imprints of immune or treatment adaptation in the donor can severely modulate viral load in the recipient [47, 5]. The reversion of transmitted ‘deleterious’ escape variants can be hindered by local fitness optima [48], potentially allowing for a transitory effect on population-level viral load [5, 42].

A heterogeneous, rapidly evolving within-host population stimulates competition between HIV-1 strains during the course of infection that will maximise viral fitness, and this short-sighted evolution results in increased virulence [24, 23]. To what extent increased replicative capacity results in higher HIV-1 virulence is still controversial [35, 19, 23, 49], given the moderate correlation between viral load and CD4 count, and other factors likely also influence virulence [42, 35]. Additionally, recent studies on the heritability of virulence show that the virus genotype affects HIV-1 virulence both via viral load and viral load-independent mechanisms [50, 49, 35, 51, 49, 52, 15, 46].

## 3 Selective bottlenecks in HIV-1 transmission

The extent to which within-host evolution has an influence on between-host transmission further depends on the social dynamics of the host population and the biological processes defining the transmission event [53, 42, 54, 27, 41, 5, 55].

The time between transmission events is particularly of interest [23]. Transmission events in early infection limit the time window for adaptation to the host and will minimize the impact of within-host evolution. Longer time intervals are accompanied by increasing viral evolution, which could modulate the transmission fitness of the viral population. Although infectiousness is known to peak during acute infection, the relative contribution of transmissions during this phase over later disease stages varies between studies [56, 57, 58, 59]. One study recently questioned established estimates of elevated infectivity in acute phase, due to the presence of confounding factors such as risk behaviour [58]. Documented transmission of drug resistance and preliminary findings on TasP effectiveness indicate that HIV-1 is transmitted beyond acute infection to some extent [57, 60]. An accurate timing of transmissions will be highly relevant for TasP policies as their efficacy critically depends on early diagnosis of HIV-1 infections, which is generally difficult to achieve.

Another important parameter relate to the diverse immunological and physical barriers in both donor and recipient that result in a low probability of transmission per contact act [61, 57]. The existence of a strong genetic bottleneck is illustrated by a genetically homogeneous virus population present in recipients shortly after infection, compared to extensively diversified populations in donors [59]. Systemic infection is mainly established from the propagation of a single variant, the transmitted/founder (T/F) virus, although estimates of multiple variant transmissions differ between transmission modes and studies [62, 55, 27, 63]. It remains unclear whether this bottleneck acts immediately at the time of transmission or in the earliest stages of infection. Another question not fully answered to date is whether particular viral traits are selectively advantageous and determine which variants survive the bottleneck, or whether successfully transmitted variants are to a considerable extent determined by stochastic effects [64]. However, the transmission process most likely involves both fitness-driven and stochastic bottlenecks in donor and recipient [59]. Comparative studies on the genetic variability of viral population in recipients versus donors have revealed that specific strains are preferentially transmitted. Moreover, analyses on T/F viruses have identified genetic, immunological and phenotypic signatures that are associated with increased transmission success [64, 42, 5, 35, 27, 53, 65].

Most well-known examples described comprise signatures of the HIV-1 envelope protein. Shorter variable loop lengths and less N-linked glycosylation sites facilitate transmission but at the cost of increased antibody recognition in the new host [64]. Additionally, T/F viruses predominantly use the CCR5 co-receptor for cell entry, which could suggest that CXCR4-utilizing viruses are associated with reduced transmissibility, although it has been speculated that CXCR4 variants are co-transmitted but outcompeted by CCR5 variants upon transmission [51, 26, 66]. Non-envelope signatures have also been associated with transmission efficiency, although less pronounced and still highly debated [5, 26, 67]. It has been postulated that the genetic bottleneck imposes a transmission selection bias that favours viruses with higher transmission potential [5, 53, 65], and this bias differs in strength according to the transmission risk associated with the specific barrier [53]. For example, the use of microbicide gels could select for viruses with higher transmission fitness while genital inflammation exerts an opposite effect since it facilitates transmission [53]. Similarly, the use of antivirals for HIV-1 negative individuals at high risk for infection, known as Pre-exposure prophylaxis (PrEP), also affects this transmission barrier. PrEP has been highly effective in reducing HIV-1 incidence [68], although a PreP failure was recently documented despite good adherence [69]. This raises important concerns with respect to the propagation of transmitted drug resistance and the amplification of virulence evolution [70, 71].

Furthermore, transmission fitness has been linked to the similarity of the T/F virus to the HIV-1 (subtype) consensus sequence among infected individuals [72]. The preferential transmission of viruses more similar to the ancestral donor strain suggests that viruses more related to the previously transmitted variant have a higher transmission efficiency [65].

In summary, while a consensus exists that selection is acting to some extent during the transmission event, favouring particular viral variants, it is not yet clear how the different factors interplay with each other.

## 4 Paradox between HIV-1 evolution at host and population level

HIV-1 evolution is not only shaped by constraints within the host and during a single transmission process but also at the population level. Increased intensity of viral competition in the infected individual results in decreased fitness of the viral population at the epidemiological level [23].

Consequently, the evolution of HIV-1 virulence has been widely studied for its impact on transmission dynamics and epidemic spread [23, 49]. More specifically, a highly virulent virus is associated with a higher viral load and increased infectiousness, but the time window for its transmission is reduced due to accelerated progression to AIDS-related death [49]. On the contrary, a low virulent virus will be less harmful to the host but the longer duration of infection provides more opportunities to establish onward transmission, although with a lower probability of transmission per contact [23].

In this context, while 20% of spVL variation can be explained by host genetics and demography [73, 18], a strong correlation of spVL between transmission pairs indicates that spVL variation can be significantly attributed to the viral genotype [74], with spVL heritability estimates averaging around 30% [50, 75, 18, 76]. A consensus on the transmitted viral characteristics determining viral load heritability and their strength has still not been established [19, 20, 77].

Observations that spVL varies between individuals, influences between-host fitness and is heritableimply that this viral trait is subject to selection at population level [24, 25]. Indeed, it has been suggested that viral genotypes with intermediate virulence are naturally selected by transmission, evolving to a spVL that should reflect the optimal balance between infectiousness and duration of infection [25, 52, 78, 79]. The quantification of this transmission potential has been validated using mathematical models and observations of population viral load distributions that are centered around the value predicted to optimize transmissibility [25, 52, 75, 22]. The existence of such an trade-off between virulence and transmissibility at population level is remarkable given the shortsighted nature of HIV-1 within-host evolution [24, 22, 23, 80].

Mathematical models suggest that HIV-1 within-host adaptation contributes more to virus genetic variability than adaptation to reach a higher transmissibility, despite observations of population level adaptation [81]. Different theories have been proposed to reconcile observed adaptation patterns at the within-and between-host level, explaining how short-sighted evolution is bypassed and viruses with high transmission potential are favoured [78, 19, 20, 45, 82, 80, 83]. Fast replicating viruses might not achieve expected high prevalence due to higher levels of immune selective pressure against them [19, 5]. An alternative mechanism is that factors such as virus-induced cell activation, acting at the systemic level, affect spVL, but are selectively neutral or nearly neutral [84, 20, 19]. The preferential transmission of ancestral viruses with an inherent transmission advantage, either early in infection or transiently released from long-lived latent reservoirs, could also minimize the impact of adaptation to the host [45, 19].

## 5 Implications for transmission analyses

The inference of HIV-1 transmission dynamics has become increasingly important to design efficient public health interventions [10]. Most methods identify clusters in virus genetic data either by comparing genetic distances directly or by interpreting phylogenetic trees [10]. High evolutionary rates of HIV-1 allow researchers to study virus spread even for short timescales, but discrepancies between population-level and within-host evolution confound the inference of epidemiological parameters of HIV-1 spread [10, 85, 86, 87]. The evolutionary history of HIV-1 can be reconstructed by linking phylogenetic methods with epidemiological models, thereby revealing dates and ordering of transmission events. However, a mismatch between actual transmission histories and time-scaled between-host phylogenies can occur when within-host evolution in the donor is neglected [85, 87, 86].

A split in a phylogenetic tree does not directly correspond to a transmission event but also reflects the sampling of (long-persisting) viral lineages in the donor at the time of transmission [85]. The presence of a pre-transmission interval, the time difference between the origin of the transmitted virus lineage and the actual transmission, pushes estimated times of transmission events backwards in time compared to the actual time of transmission [85]. The impact of this interval depends on the amount of viral diversity accumulated in the donor and hence on the duration of infection. Besides this bias on estimates of time of transmission, the order in which nodes occur in an estimated phylogeny can also be confounded by within-host dynamics and sampling times (Figure 3). An example is incomplete lineage sorting which occurs when a single donor infects multiple individuals, thereby transmitting lineages from different time points [85, 87, 86]. The extent of disordering of tree nodes is affected by viral diversity in the donor and by the length of the time intervals between the separate events [85].

**Figure 3:**
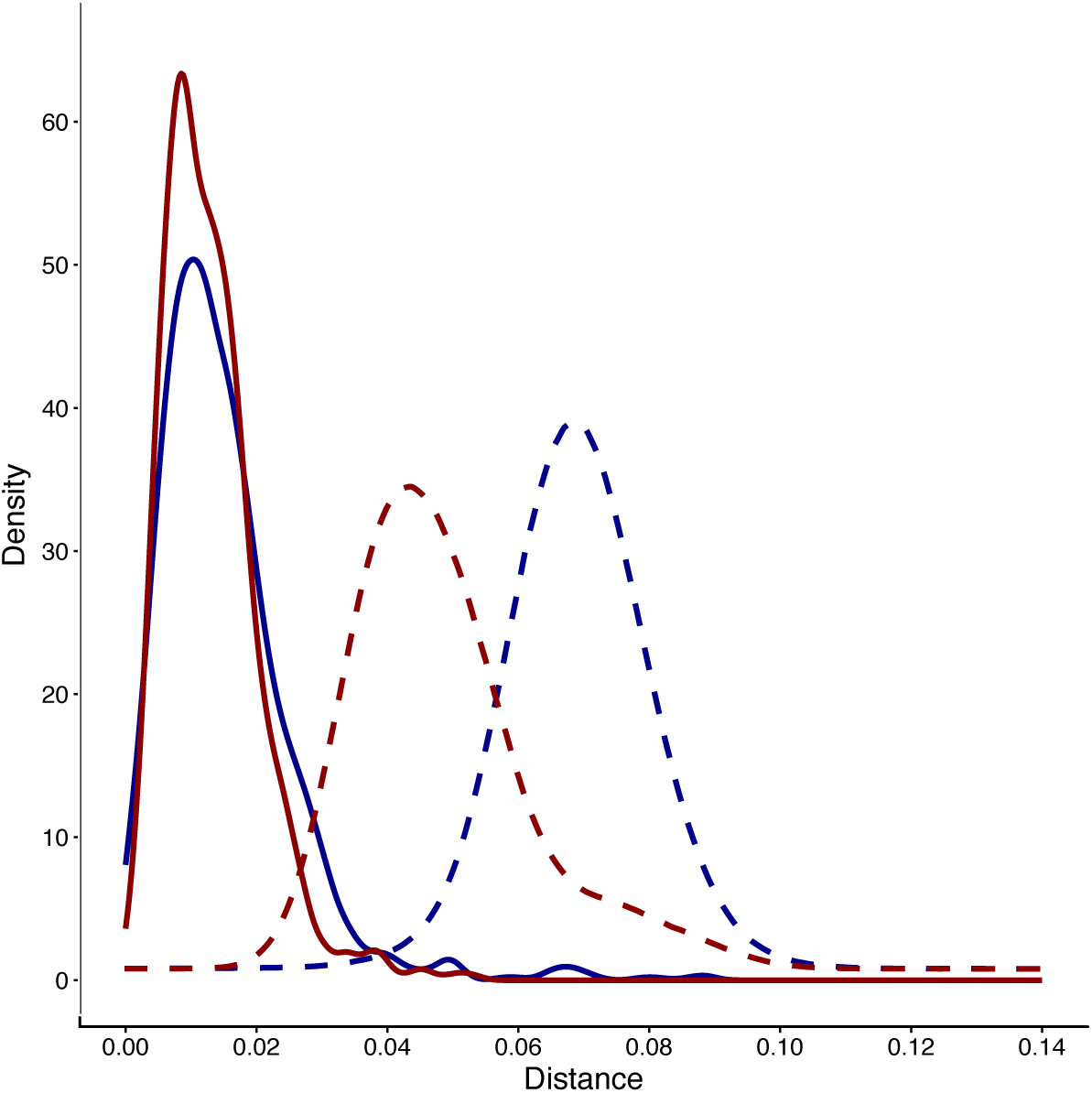
In this figure, we demonstrate how the inter-patient distance distribution can differ between HIV-1 sub-epidemics. We consider the Portuguese HIV-1 epidemic, as this epidemic consists of two parallel sub-epidemics that can be easily discerned by their subtype (i.e., subtype B and subtype G) [91]. We extracted 6045 subtype B and 4311 subtype G sequences from 7977 number of patients the Egas Moniz HIV-1 resistance RegaDB database [92, 93] and inferred the subtype of the sequences using the Rega HIV-1 typing tool [94]. For both subtypes the intra-host and inter-host pairwise (Tamura-Nei) distances were determined [95]. We show distance distributions for subtype B (red density curve) and subtype G (blue density curve). The solid lines represent the intra-patient distance distribution’s density, the dashed lines represent the density of the inter-patient distance distribution.

Moreover, the identification of transmission clusters based on genetic distance typically relies on a predefined threshold [88]. This threshold should maximally separate within-host from between-host distance distributions. Figure 4 shows that genetic distance distributions can differ between sub-epidemics, indicating that such thresholds should be specifically tailored to the analysed sub-epidemic.

**Figure 4:**
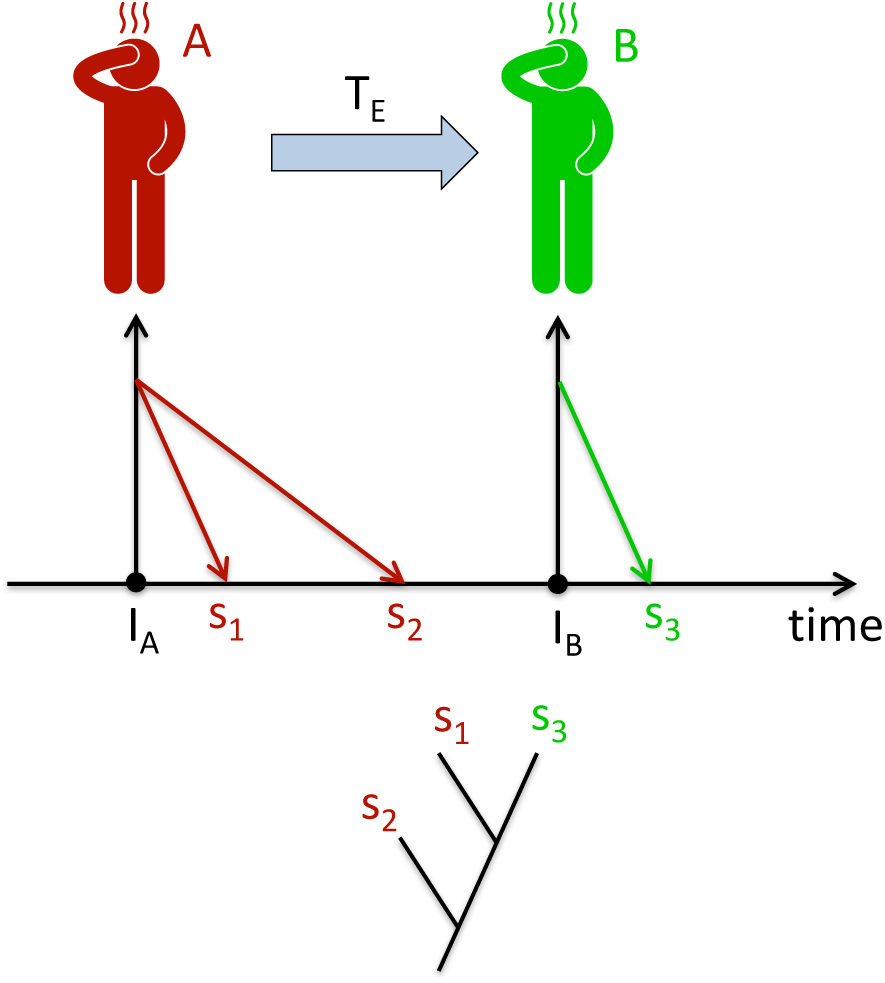
We show a schematic visualization of the the transmission event T_E_, where individual A infects individual B at time I_B_. We consider 2 samples from individual A: s_1_ was obtained shortly after individual A was infected (i.e., time I_A_) and sample S_2_ was obtained close to time of transmission event T_E_. A third sample S3 was obtained from individual B, shortly after this individual was infected. The phylogenetic tree, as inferred from the genetic sequences extracted from sample s_1_, s_2_ and S_3_, reveals that s_1_ and S_3_ cluster more closely together than s_1_ and s_2_.

More realistic phylodynamic models are acknowledging therole of within-host diversification and will improve the inference of transmission dynamics [86, 87]. Increased knowledge on the impact of within-host evolution also benefits fine-grained epidemiological models, which are essential to calibrate complex prevention strategies such as test-and-treat protocols [89].

## 6 Conclusions

While multiple studies illustrate that HIV-1 is adapting to maximize transmissibility, the consequences of within-host evolutionary processes on HIV-1 dynamics at the epidemiological level are not yet fully understood. Future studies will bring insights into which variants are transmitted and the dynamics of their global spread, which will ultimately inform prevention and vaccine strategies which host or virus populations to target and at which time.

## 7 Acknowledgements

Kristof Theys is funded by a postdoctoral grant of the Fonds Wetenschappelijk Onderzoek Vlaanderen (FWO). Pieter Libin is funded by a doctoral grant of the Research Foundation - Flanders (FWO). This work has been funded by the Fonds voor Wetenschappelijk Onderzoek Flanders (FWO) grant G.0692.14, and G.0611.09N, Fundao para a Cincia e Tecnologia (FCT) through funds to GHTM-UID/Multi/04413/2013, the VIROGENESIS project that receives funding from the European Unions Horizon 2020 research and innovation program under grant agreement No 634650, by the BEST HOPE project (Bio-Molecular and Epidemiological Surveillance of HIV Transmitted Drug Resistance, Hepatitis Co-Infections and Ongoing Transmission Patterns in Europe’) (project 35 funded through HIVERA: Harmonizing Integrating 450 Vitalizing European Research on HIV/Aids, grant 249697), by the MigrantHIV project (financed by FCT: PTDC/DTP-EPI/7066/2014) and by the Gilead Genese programme.

## References

1. R. Shankarappa, J. B Margolick, S. J. Gange, A. G Rodrigo, D. Upchurch, H. Farzadegan, P. Gupta, C. R Rinaldo, G. H. Learn, X. He, X. L Huang, J. I. Mullins, Consistent viral evolutionary changes associated with the progression of human immunodeficiency virus type 1 infection, J. Virol. 73 (12) (1999) 10489–10502.

2. P. Lemey, A. Rambaut, O. G Pybus, HIV evolutionary dynamics within and among hosts, AIDS Rev 8 (3) (2006) 125–140.

3. P. Lemey, S. L. Kosakovsky Pond, A. J Drummond, O. G. Pybus, B. Shapiro, H. Barroso, N. Taveira, A. Rambaut, Synonymous substitution rates predict HIV disease progression as a result of underlying replication dynamics, PLoS Comput. Biol. 3 (2) (2007) e29.

4. P. A. Volberding, S. G Deeks, Antiretroviral therapy and management of HIV infection, Lancet 376 (9734) (2010) 49–62.

5. J. M. Carlson, M. Schaefer, D. C Monaco, R. Batorsky, D. T Claiborne, J. Prince, M. J Deymier, Z. S. Ende, N. R Klatt, C. E. DeZiel, T. H Lin, J. Peng, A. M Seese, R. Shapiro, J. Frater, T. Ndung’u, J. Tang, P. Goepfert, J. Gilmour, M. A Price, W. Kilembe, D. Heckerman, P. J Goulder, T. M. Allen, S. Allen, E. Hunter, HIV transmission. Selection bias at the heterosexual HIV-1 transmission bottleneck, Science 345 (6193) (2014) 1254031.

6. H. Wang, M. Naghavi, C. Allen, R. M Barber, Z. A. Bhutta, A. Carter, D. C Casey, F. J. Charlson, A. Z Chen, M. M. Coates, M. Coggeshall, L. Dandona, D. J Dicker, H. E. Erskine, A. J Ferrari, Global, regional, and national life expectancy, all-cause mortality, and cause-specific mortality for 249 causes of death, 1980-2015: a systematic analysis for the Global Burden of Disease Study 2015, Lancet 388 (10053) (2016) 1459–1544.

7. A. Trickey, M. T May, J. J. Vehreschild, N. Obel, M. J Gill, H. M. Crane, C. Boesecke, S. Patterson, S. Grabar, C. Cazanave, M. Cavassini, L. Shepherd, A. D Monforte, A. van Sighem, M. Saag, F. Lampe, V. Hernando, M. Montero, R. Zangerle, A. C Justice, T. Sterling, S. M Ingle, J. A. C Sterne, Survival of HIV-positive patients starting antiretroviral therapy between 1996 and 2013: a collaborative analysis of cohort studies, Lancet HIV 4 (8) (2017) e349–e356.

8. A. J. McMichael, P. Borrow, G. D Tomaras, N. Goonetilleke, B. F Haynes, The immune response during acute HIV-1 infection: clues for vaccine development, Nat. Rev. Immunol. 10 (1) (2010) 11–23.

9. M. S. Cohen, M. K Smith, K. E. Muessig, T. B Hallett, K. A. Powers, A. D Kashuba, Antiretroviral treatment of HIV-1 prevents transmission of HIV-1: where do we go from here?, Lancet 382 (9903) (2013) 1515–1524.

10. A. F. Y Poon, Impacts and shortcomings of genetic clustering methods for infectious disease outbreaks, Virus Evolution 2 (2) (2016) vew031. arXiv:/oup/backfile/content_public/journal/ve/2/2/10.1093_ve_vew031/1/vew031.pdf, doi:10.1093/ve/vew031. URL +http://dx.doi.org/10.1093/ve/vew031

11. A. Pharris, C. Quinten, L. Tavoschi, G. Spiteri, A. J. Amato-Gauci, D. Schmid, A. Sasse, D. Van Beckhoven, T. Varleva, T. Nemeth Blazic, M. Koliou, L. Hadjihannas, M. Maly, S. Cowan, K. Ruutel, K. Liitsola, M. Salminen, F. Cazein, J. Pillonel, F. Lot, B. Gunsenheimer-Bartmeyer, G. Nikolopoulos, D. Paraskeva, M. Dudas, H. Briem, G. Sigmundsdottir, D. Igoe, K. O’Donnell, D. O’Flanagan, B. Suligoi, S. Konova, S. Erne, I. Caplinskiene, J. C Schmit, J. M. Melillo, T. Melillo, E. Op de Coul, H. Blystad, M. Rosinska, A. Diniz, M. Mardarescu, P. Truska, I. Klavs, M. Diez, M. Axelsson, V. Delpech, Trends in HIV surveillance data in the EU/EEA, 2005 to 2014: new HIV diagnoses still increasing in men who have sex with men, Euro Surveill. 20 (47).

12. M. Plazy, K. E Farouki, C. Iwuji, N. Okesola, J. Orne-Gliemann, J. Lar-marange, F. Lert, M. L Newell, F. Dabis, R. Dray-Spira, Access to HIV care in the context of universal test and treat: challenges within the ANRS 12249 TasP cluster-randomized trial in rural South Africa, J Int AIDS Soc 19 (1) (2016) 20913.

13. E. Raffetti, M. C Postorino, F. Castelli, S. Casari, F. Castelnuovo, F. Mag-giolo, E. Di Filippo, A. D’Avino, A. Gori, N. Ladisa, M. Di Pietro, L. Sigh-inolfi, F. Zacchi, C. Torti, The risk of late or advanced presentation of HIV infected patients is still high, associated factors evolve but impact on overall mortality is vanishing over calendar years: results from the Italian MASTER Cohort, BMC Public Health 16 (1) (2016) 878.

14. G. Li, S. Piampongsant, N. R Faria, A. Voet, A. C. Pineda-Pena, R. Khouri, P. Lemey, A. M Vandamme, K. Theys, An integrated map of HIV genome-wide variation from a population perspective, Retrovirology 12 (2015) 18.

15. J. Hemelaar, Implications of HIV diversity for the HIV-1 pandemic, J. Infect. 66 (5) (2013) 391–400.

16. N. R. Faria, A. Rambaut, M. A Suchard, G. Baele, T. Bedford, M. J Ward, A. J. Tatem, J. D Sousa, N. Arinaminpathy, J. Pepin, D. Posada, M. Peeters, O. G Pybus, P. Lemey, HIV epidemiology. The early spread and epidemic ignition of HIV-1 in human populations, Science 346 (6205) (2014) 56–61.

17. A. C. Pineda-Pena, J. Varanda, J. D Sousa, K. Theys, I. Bartolo, T. Leitner, N. Taveira, A. M Vandamme, A. B. Abecasis, On the contribution of Angola to the initial spread of HIV-1, surname>Infect. Genet. Evol. 46 (2016) 219–222.

18. I. Bartha, P. J. McLaren, C. Brumme, R. Harrigan, A. Telenti, J. Fellay, Estimating the Respective Contributions of Human and Viral Genetic Variation to HIV Control, PLoS Comput. Biol. 13 (2) (2017) e1005339.

19. C. Fraser, K. Lythgoe, G. E Leventhal, G. Shirreff, T. D Hollingsworth, S. Alizon, S. Bonhoeffer, Virulence and pathogenesis of HIV-1 infection: an evolutionary perspective, Science 343 (6177) (2014) 1243727.

20. S. Bonhoeffer, C. Fraser, G. E Leventhal, High heritability is compatible with the broad distribution of set point viral load in HIV carriers, PLoS Pathog. 11 (2) (2015) e1004634.

21. A. Rambaut, D. L Robertson, O. G. Pybus, M. Peeters, E. C Holmes, Human immunodeficiency virus. Phylogeny and the origin of HIV-1, Nature 410 (6832) (2001) 1047–1048.

22. K. A. Lythgoe, L. Pellis, C. Fraser, Is HIV short-sighted? Insights from a multistrain nested model, Evolution 67 (10) (2013) 2769–2782.

23. K. A. Lythgoe, A. Gardner, O. G Pybus, J. Grove, Short-Sighted Virus Evolution and a Germline Hypothesis for Chronic Viral Infections, Trends Microbiol. 25 (5) (2017) 336–348.

24. B. R. Levin, J. J Bull, Short-sighted evolution and the virulence of pathogenic microorganisms, Trends Microbiol. 2 (3) (1994) 76–81.

25. C. Fraser, T. D Hollingsworth, R. Chapman, F. de Wolf, W. P. Hanage, Variation in HIV-1 set-point viral load: epidemiological analysis and an evolutionary hypothesis, Proc. Natl. Acad. Sci. U.S.A. 104 (44) (2007) 17441–17446.

26. C. S. Oberle, B. Joos, P. Rusert, N. K Campbell, D. Beauparlant, H. Kuster, J. Weber, C. D Schenkel, A. U. Scherrer, C. Magnus, R. Kouyos, P. Rieder, B. Niederost, D. L Braun, J. Pavlovic, J. Boni, S. Yerly, T. Klimkait, V. Aubert, A. Trkola, K. J Metzner, H. F. Gunthard, V. Aubert, M. Battegay, E. Bernasconi, J. Boni, D. Braun, H. Bucher, C. Burton-Jeangros, A. Calmy, M. Cavassini, G. Dollenmaier, M. Egger, L. Elzi, J. Fehr, J. Fellay, H. Furrer, C. Fux, M. Gorgievski, H. Gunthard, D. Haerry, B. Hasse, H. Hirsch, M. Hoffmann, I. Hosli, C. Kahlert, L. Kaiser, O. Keiser, T. Klimkait, R. Kouyos, H. Kovari, B. Ledergerber, G. Martinetti, B. Martinez de Tejada, C. Marzolini, K. Metzner, N. Muller, D. Nadal, D. Nicca, G. Pantaleo, A. Rauch, S. Regenass, C. Rudin, F. Schoni-Affolter, P. Schmid, R. Speck, M. Stockle, P. Tarr, A. Trkola, P. Vernazza, R. Weber, S. Yerly, Tracing HIV-1 transmission: envelope traits of HIV-1 transmitter and recipient pairs, Retrovirology 13 (1) (2016) 62.

27. D. C. Tully, C. B Ogilvie, R. E. Batorsky, D. J Bean, K. A. Power, M. Ghebremichael, H. E Bedard, A. D. Gladden, A. M Seese, M. A. Amero, K. Lane, G. McGrath, S. B Bazner, J. Tinsley, N. J Lennon, M. R. Henn, Z. L Brumme, P. J. Norris, E. S Rosenberg, K. H. Mayer, H. Jessen, S. L. Kosakovsky Pond, B. D Walker, M. Altfeld, J. M Carlson, T. M. Allen, Differences in the Selection Bottleneck between Modes of Sexual Transmission Influence the Genetic Composition of the HIV-1 Founder Virus, PLoS Pathog. 12 (5) (2016) e1005619.

28. R. B. Geskus, M. Prins, J. B Hubert, F. Miedema, B. Berkhout, C. Rouzioux, J. F Delfraissy, L. Meyer, The HIV RNA setpoint theory revisited, Retrovirology 4 (2007) 65.

29. G. Doitsh, W. C Greene, Dissecting How CD4 T Cells Are Lost During HIV Infection, Cell Host Microbe 19 (3) (2016) 280–291.

30. N. Blaser, C. Wettstein, J. Estill, L. S Vizcaya, G. Wandeler, M. Egger, O. Keiser, Impact of viral load and the duration of primary infection on HIV transmission: systematic review and meta-analysis, AIDS 28 (7) (2014) 1021–1029.

31. R. H. Gray, M. J Wawer, R. Brookmeyer, N. K Sewankambo, D. Serwadda, F. Wabwire-Mangen, T. Lutalo, X. Li, T. vanCott, T. C Quinn, Probability of HIV-1 transmission per coital act in monogamous, heterosexual, HIV-1-discordant couples in Rakai, Uganda, Lancet 357 (9263) (2001) 1149–1153.

32. J. W. Mellors, C. R Rinaldo, P. Gupta, R. M White, J. A. Todd, L. A Kingsley, Prognosis in HIV-1 infection predicted by the quantity of virus in plasma, Science 272 (5265) (1996) 1167–1170.

33. M. A. Brockman, Z. L Brumme, C. J. Brumme, T. Miura, J. Sela, P. C Rosato, C. M. Kadie, J. M Carlson, T. J. Markle, H. Streeck, A. D. Kelleher, M. Markowitz, H. Jessen, E. Rosenberg, M. Altfeld, P. R Harrigan, D. Heckerman, B. D Walker, T. M. Allen, Early selection in Gag by protective HLA alleles contributes to reduced HIV-1 replication capacity that may be largely compensated for in chronic infection, J. Virol. 84 (22) (2010) 11937–11949.

34. J. K. Wright, Z. L Brumme, J. M. Carlson, D. Heckerman, C. M Kadie, C. J. Brumme, B. Wang, E. Losina, T. Miura, F. Chonco, M. van der Stok, Z. Mncube, K. Bishop, P. J Goulder, B. D. Walker, M. A Brockman, T. Ndung’u, Gag-protease-mediated replication capacity in HIV-1 subtype C chronic infection: associations with HLA type and clinical parameters, J. Virol. 84 (20) (2010) 10820–10831.

35. D. T. Claiborne, J. L Prince, E. Scully, G. Macharia, L. Micci, B. Lawson, J. Kopycinski, M. J Deymier, T. H. Vanderford, K. Nganou-Makamdop, Z. Ende, K. Brooks, J. Tang, T. Yu, S. Lakhi, W. Kilembe, G. Silvestri, D. Douek, P. A Goepfert, M. A. Price, S. A Allen, M. Paiardini, M. Altfeld, J. Gilmour, E. Hunter, Replicative fitness of transmitted HIV-1 drives acute immune activation, proviral load in memory CD4+ T cells, and disease progression, Proc. Natl. Acad. Sci. U.S.A. 112 (12) (2015) E1480–1489.

36. L. Yue, K. J Pfafferott, J. Baalwa, K. Conrod, C. C Dong, C. Chui, R. Rong, D. T Claiborne, J. L. Prince, J. Tang, R. M Ribeiro, E. Cormier, B. H Hahn, A. S. Perelson, G. M Shaw, E. Karita, J. Gilmour, P. Goepfert, C. A Derdeyn, S. A. Allen, P. Borrow, E. Hunter, Transmitted virus fitness and host T cell responses collectively define divergent infection outcomes in two HIV-1 recipients, PLoS Pathog. 11 (1) (2015) e1004565.

37. F. Bielejec, G. Baele, A. G Rodrigo, M. A. Suchard, P. Lemey, Identifying predictors of time-inhomogeneous viral evolutionary processes, Virus Evol 2 (2) (2016) vew023.

38. A. S. Olabode, S. M Kandathil, S. C. Lovell, D. L Robertson, Adaptive HIV-1 evolutionary trajectories are constrained by protein stability, Virus Evol 3 (2) (2017) vex019.

39. E. S. Daar, K. L Kesler, T. Wrin, C. J Petropoulo, M. Bates, A. Lail, N. S Hellmann, E. Gomperts, S. Donfield, HIV-1 pol replication capacity predicts disease progression, AIDS 19 (9) (2005) 871–877.

40. R. D. Kouyos, V. von Wyl, T. Hinkley, C. J Petropoulos, M. Haddad, J. M Whitcomb, J. Boni, S. Yerly, C. Cellerai, T. Klimkait, H. F Gunthard, S. Bonhoeffer, Assessing predicted HIV-1 replicative capacity in a clinical setting, PLoS Pathog. 7 (11) (2011) e1002321.

41. J. M. Carlson, A. Q Le, A. Shahid, Z. L Brumme, HIV-1 adaptation to HLA: a window into virus-host immune interactions, Trends Microbiol. 23 (4) (2015) 212–224.

42. J. M. Carlson, V. Y Du, N. Pfeifer, A. Bansal, V. Y Tan, K. Power, C. J Brumme, A. Kreimer, C. E. DeZiel, N. Fusi, M. Schaefer, M. A Brockman, J. Gilmour, M. A Price, W. Kilembe, R. Haubrich, M. John, S. Mallal, R. Shapiro, J. Frater, P. R Harrigan, T. Ndung’u, S. Allen, D. Heckerman, J. Sidney, T. M Allen, P. J. Goulder, Z. L Brumme, E. Hunter, P. A Goepfert, Impact of pre-adapted HIV transmission, Nat. Med. 22 (6) (2016) 606–613.

43. R. Winand, K. Theys, M. Eusebio, J. Aerts, R. J Camacho, P. Gomes, M. A Suchard, A. M. Vandamme, A. B Abecasis, Assessing transmissibility of HIV-1 drug resistance mutations from treated and from drug-naive individuals, AIDS 29 (15) (2015) 2045–2052.

44. K. Theys, K. Deforche, J. Vercauteren, P. Libin, D. A. van de Vijver, J. Albert, B. Asjo, C. Balotta, M. Bruckova, R. J Camacho, B. Clotet, S. Coughlan, Z. Grossman, O. Hamouda, A. Horban, K. Korn, L. G Kostrikis, C. Kucherer, C. Nielsen, D. Paraskevis, M. Poljak, E. Puchhammer-Stockl, C. Riva, L. Ruiz, K. Liitsola, J. C Schmit, R. Schuurman, A. Sonnerborg, D. Stanekova, M. Stanojevic, D. Struck, K. Van Laethem, A. M Wensing, C. A. Boucher, A. M Vandamme, E. Puchhammer-Stockl, M. Sarcletti, B. Schmied, M. Geit, G. Balluch, A. Vandamme, J. Vercauteren, I. Derdelinckx, A. Sasse, M. Bogaert, H. Ceunen, A. De Roo, S. De Wit, K. Deforche, F. Echahidi, K. Fransen, J. Goffard, P. Goubau, E. Goudeseune, J. Yombi, P. Lacor, C. Liesnard, M. Moutschen, D. Pierard, R. Rens, Y. Schrooten, D. Vaira, A. Van den Heuvel, B. Van Der Gucht, M. Van Ranst, E. Van Wijngaerden, B. Vandercam, M. Vekemans, C. Verhofstede, N. Clumeck, K. Van Laethem, L. Kostrikis, I. Demetriades, I. Kousiappa, V. Demetriou, J. Hezka, M. Linka, M. Bruckova, L. Machala, C. Nielsen, L. Jrgensen, J. Gerstoft, L. Mathiesen, C. Pedersen, H. Nielsen, A. Laursen, B. Kvinesdal, K. Liitsola, M. Ristola, J. Suni, J. Sutinen, C. Kucherer, T. Berg, P. Braun, G. Poggensee, M. Daumer, J. Eberle, O. Hamouda, H. Heiken, R. Kaiser, H. Knechten, H. Muller, S. Neifer, B. Schmidt, H. Walter, B. Gunsenheimer-Bartmeyer, T. Harrer, A. Hatzakis, D. Paraskevis, E. Magiorkinis, E. Hatzitheodorou, C. Issaris, C. Haida, A. Zavitsanou, G. Magiorkinis, M. Lazanas, M. Chini, N. Magafas, N. Tsogas, V. Paparizos, S. Kourkounti, A. Antoniadou, A. Papadopoulos, P. Panagopoulos, G. Poulakou, V. Sakka, G. Chryssos, S. Drimis, P. Gargalianos, M. Lelekis, G. Xilomenos, M. Psichogiou, G. Daikos, G. Panos, G. Haratsis, T. Kordossis, A. Kontos, G. Koratzanis, M. Theodoridou, G. Mostrou, V. Spoulou, C. De Gascun, C. Byrne, M. Duffy, C. Bergin, D. Reidy, G. Farrell, J. Lambert, E. O’Connor, A. Rochford, J. Low, P. Coakely, S. Coughlan, Z. Grossman, I. Levi, D. Chemtob, C. Balotta, C. Riva, C. Mussini, I. Caramma, A. Capetti, M. Colombo, C. Rossi, F. Prati, F. Tramuto, F. Vitale, M. Ciccozzi, G. Angarano, G. Rezza, J. Schmit, D. Struck, R. Hemmer, V. Arendt, T. Staub, F. Schneider, F. Roman, A. Wensing, C. Boucher, D. van de Vijver, P. van Bentum, K. Brinkman, E. de Coul, M. van der Ende, I. Hoepelman, M. van Kasteren, J. Juttmann, M. Kuipers, N. Langebeek, C. Richter, R. Santegoets, L. Schrijnders-Gudde, R. Schuurman, B. van de Ven, B. Asjo, V. Ormaasen, P. Aavitsland, A. Horban, J. Stanczak, G. Stanczak, E. Firlag-Burkacka, A. Wiercinska-Drapalo, E. Jablonowska, E. Malolepsza, M. Leszczyszyn-Pynka, W. Szata, R. Camacho, C. Palma, F. Borges, T. Paixao, V. Duque, F. Araujo, M. Stanojevic, D. J Jevtovic, D. Salemovic, D. Stanekova, M. Habekova, M. Mokras, P. Truska, M. Poljak, D. Babic, J. Tomazic, L. Vidmar, P. Karner, B. Clotet, C. Gutierrez, C. de Mendoza, I. Erkicia, P. Domingo, X. Camino, M. Galindo, J. Blanco, M. Leal, A. Masabeu, A. Guelar, J. Llibre, N. Margall, J. Iribarren, S. Gutierrez, J. Baldovi, J. Pedreira, J. Gatell, S. Moreno, C. de Mendoza, V. Soriano, L. Ruiz, J. Albert, A. Blaxhult, A. Heidarian, A. Karlsson, K. Aperia-Peipke, I. Bergbrant, M. Gisslen, B. Svennerholm, P. Bjorkman, G. Bratt, M. Carlsson, H. Ekvall, M. Ericsson, M. Hofer, B. Johansson, A. Sonnerborg, N. Kuylenstierna, B. Ljungberg, S. Makitalo, A. Strand, S. Oberg, Treatment-associated polymorphisms in protease are significantly associated with higher viral load and lower CD4 count in newly diagnosed drug-naive HIV-1 infected patients, Retrovirology 9 (2012) 81.

45. I. M. Rouzine, A. D Weinberger, L. S. Weinberger, An evolutionary role for HIV latency in enhancing viral transmission, Cell 160 (5) (2015) 1002–1012.

46. V. Kouri, R. Khouri, Y. Aleman, Y. Abrahantes, J. Vercauteren, A. C. Pineda-Pena, K. Theys, S. Mekgens, M. Moutschen, N. Pfeifer, J. Van Weyenbergh, A. B Perez, J. Perez, L. Perez, K. Van Laethem, A. M Vandamme, CRF19cpx is an Evolutionary fit HIV-1 Variant Strongly Associated With Rapid Progression to AIDS in Cuba, EBioMedicine 2 (3) (2015) 244–254.

47. P. A. Goepfert, W. Lumm, P. Farmer, P. Matthews, A. Prendergast, J. M Carlson, C. A. Derdeyn, J. Tang, R. A Kaslow, A. Bansal, K. Yusim, D. Heckerman, J. Mulenga, S. Allen, P. J Goulder, E. Hunter, Transmission of HIV-1 Gag immune escape mutations is associated with reduced viral load in linked recipients, J. Exp. Med. 205 (5) (2008) 1009–1017.

48. R. D. Kouyos, H. F Gunthard, Editorial Commentary: The Irreversibility of HIV Drug Resistance, Clin. Infect. Dis. 61 (5) (2015) 837–839.

49. F. Bertels, A. Marzel, G. Leventhal, V. Mitov, J. Fellay, H. F. Gunthard, J. Boni, S. Yerly, T. Klimkait, V. Aubert, M. Battegay, A. Rauch, M. Cavassini, A. Calmy, E. Bernasconi, P. Schmid, A. Scherrer, V. Muller, S. Bonhoeffer, R. Kouyos, R. R Regoes, Dissecting hiv virulence: Heritability of setpoint viral load, cd4+ t cell decline and per-parasite pathogenicity, bioRxivarXiv: http://www.biorxiv.org/content/early/2017/05/22/140012.full.pdf, doi:10.1101/140012. URL http://www.biorxiv.org/content/early/2017/05/22/140012

50. S. Alizon, V. von Wyl, T. Stadler, R. D Kouyos, S. Yerly, B. Hirschel, J. Boni, C. Shah, T. Klimkait, H. Furrer, A. Rauch, P. L Vernazza, E. Bernasconi, M. Battegay, P. Burgisser, A. Telenti, H. F Gunthard, S. Bonhoeffer, Phylogenetic approach reveals that virus genotype largely determines HIV set-point viral load, PLoS Pathog. 6 (9) (2010) e1001123.

51. J. M. Baeten, B. Chohan, L. Lavreys, V. Chohan, R. S. McClelland, L. Certain, K. Mandaliya, W. Jaoko, J. Overbaugh, HIV-1 subtype D infection is associated with faster disease progression than subtype A in spite of similar plasma HIV-1 loads, J. Infect. Dis. 195 (8) (2007) 1177–1180.

52. F. Blanquart, M. K Grabowski, J. Herbeck, F. Nalugoda, D. Serwadda, M. A Eller, M. L. Robb, R. Gray, G. Kigozi, O. Laeyendecker, K. A. Lythgoe, G. Nakigozi, T. C Quinn, S. J. Reynolds, M. J Wawer, C. Fraser, A transmission-virulence evolutionary trade-off explains attenuation of HIV-1 in Uganda, Elife 5.

53. N. K. Ngandu, J. M Carlson, D. R. Chopera, N. Ndabambi, Q. Abdool Karim, S. Abdool Karim, C. Williamson, Brief Report: Selection of HIV-1 Variants With Higher Transmission Potential by 1Microbicide, J. Acquir. Immune Defic. Syndr. 76 (1) (2017) 43–47.

54. N. N. Kinloch, D. R. MacMillan, A. Q Le, L. A. Cotton, D. R Bangsberg, S. Buchbinder, M. Carrington, J. Fuchs, P. R Harrigan, B. Koblin, M. Kushel, M. Markowitz, K. Mayer, M. J Milloy, M. T. Schechter, T. Wagner, B. D Walker, J. M. Carlson, A. F Poon, Z. L. Brumme, Population-Level Immune-Mediated Adaptation in HIV-1 Polymerase during the North American Epidemic, J. Virol. 90 (3) (2015) 1244–1258.

55. S. M. Kariuki, P. Selhorst, K. K Arien, J. R. Dorfman, The HIV-1 transmission bottleneck, Retrovirology 14 (1) (2017) 22.

56. C. Gay, O. Dibben, J. A Anderson, A. Stacey, A. J Mayo, P. J. Norris, J. D Kuruc, J. F. Salazar-Gonzalez, H. Li, B. F Keele, C. Hicks, D. Margolis, G. Ferrari, B. Haynes, R. Swanstrom, G. M Shaw, B. H. Hahn, J. J Eron, P. Borrow, M. S Cohen, Cross-sectional detection of acute HIV infection: timing of transmission, inflammation and antiretroviral therapy, PLoS ONE 6 (5) (2011) e19617.

57. M. S. Cohen, G. M Shaw, A. J. McMichael, B. F Haynes, Acute HIV-1 Infection, N. Engl. J. Med. 364 (20) (2011) 1943–1954.

58. S. E. Bellan, J. Dushoff, A. P Galvani, L. A. Meyers, Reassessment of HIV-1 acute phase infectivity: accounting for heterogeneity and study design with simulated cohorts, PLoS Med. 12 (3) (2015) e1001801.

59. S. B. Joseph, R. Swanstrom, A. D Kashuba, M. S. Cohen, Bottlenecks in HIV-1 transmission: insights from the study of founder viruses, Nat. Rev. Microbiol. 13 (7) (2015) 414–425.

60. A. J. Rodger, V. Cambiano, T. Bruun, P. Vernazza, S. Collins, J. van Lunzen, G. M Corbelli, V. Estrada, A. M Geretti, A. Beloukas, D. Asboe, P. Viciana, F. Gutierrez, B. Clotet, C. Pradier, J. Gerstoft, R. Weber, K. Westling, G. Wandeler, J. M Prins, A. Rieger, M. Stoeckle, T. Kummerle, T. Bini, A. Ammassari, R. Gilson, I. Krznaric, M. Ristola, R. Zangerle, P. Handberg, A. Antela, S. Allan, A. N Phillips, J. Lundgren, V. Pompeyo, M. Trastoy, R. Palacio, F. Gutierrez, M. Masia, S. Padilla, C. Robledano, B. Clotet, P. Coll, J. Pena, V. Estrada, M. Rodrigo, E. Santiago, A. Rivero, A. Antela, E. Losada, C. Lires, A. Aguilera, J. Gatell, J. Guerrero, F. Dronda, V. Soriano, D. Asboe, N. Nwokolo, J. Sewell, R. Gilson, N. Esteban, S. McNamara, A. Rodger, K. Sturgeon, M. Gompels, L. Jennings, S. Allan, C. Leen, S. Morris, M. Brady, L. Campbell, M. Fisher, J. Dhar, R. O’Connell, D. White, J. Fox, S. Fidler, P. Stanley, U. Natarajan, M. Ghanem, J. Ainsworth, A. Waters, E. Wilkins, J. Minton, J. Calderwood, H. Patel, M. Lascar, J. Lunzen, T. Kummerle, G. Fatkenheuer, E. Rund, C. Lehmann, I. Krznaric, P. Ingiliz, J. Motsch, A. Baumgarten, J. Bogner, N. Brockmeyer, H. Stellbrink, H. Jessen, J. Rockstroh, M. Stoeckle, M. Battegay, R. Weber, C. Grube, D. Braun, H. Gunthard, G. Wandeler, H. Furrer, T. Konrad, A. Rauch, P. Vernazza, M. Rasi, E. Bernasconi, P. Tarr, J. Gerstoft, T. Quist, P. Handberg, B. Clausen, L. Mathiesen, S. Oestergaard, S. Stenvang, M. Ristola, P. Kivela, K. Westling, E. Frisen, A. Blaxhult, G. Cortney, N. Clumeck, L. Vandekerckhove, J. Prins, K. Brinkman, D. Verhagen, A. Eeden, C. Pradier, J. Durant, M. Serini, S. Breaud, F. Raffi, G. Pialoux, M. Ohayon, V. Coquelin, A. Rieger, V. Touzeau-Roemer, R. Zangerle, M. Kitchen, M. Gisinger, M. Sarcletti, M. Geit, T. Bini, L. Comi, A. Pandolfo, E. Suardi, A. Ammassari, P. Pierro, G. Carli, N. Orchi, M. Celesia, C. Mussini, A. Biagio, N. Janerio, Sexual Activity Without Condoms and Risk of HIV Transmission in Serodifferent Couples When the HIV-Positive Partner Is Using Suppressive Antiretroviral Therapy, JAMA 316 (2) (2016) 171–181.

61. P. Patel, C. B Borkowf, J. T. Brooks, A. Lasry, A. Lansky, J. Mermin, Estimating per-act HIV transmission risk: a systematic review, AIDS 28 (10) (2014) 1509–1519.

62. B. F. Keele, E. E Giorgi, J. F. Salazar-Gonzalez, J. M Decker, K. T. Pham, M. G Salazar, C. Sun, T. Grayson, S. Wang, H. Li, X. Wei, C. Jiang, J. L Kirchherr, F. Gao, J. A Anderson, L. H. Ping, R. Swanstrom, G. D Tomaras, W. A. Blattner, P. A Goepfert, J. M. Kilby, M. S Saag, E. L. Delwart, M. P Busch, M. S. Cohen, D. C Montefiori, B. F. Haynes, B. Gaschen, G. S Athreya, H. Y. Lee, N. Wood, C. Seoighe, A. S Perelson, T. Bhattacharya, B. T Korber, B. H. Hahn, G. M Shaw, Identification and characterization of transmitted and early founder virus envelopes in primary HIV-1 infection, Proc. Natl. Acad. Sci. U.S.A. 105 (21) (2008) 7552–7557.

63. G. H. Kijak, E. Sanders-Buell, A. L Chenine, M. A. Eller, N. Goonetilleke, R. Thomas, S. Leviyang, E. A Harbolick, M. Bose, P. Pham, C. Oropeza, K. Poltavee, A. M. O’Sullivan, E. Billings, M. Merbah, M. C Costanzo, J. A. Warren, B. Slike, H. Li, K. K Peachman, W. Fischer, F. Gao, C. Cicala, J. Arthos, L. A Eller, R. J. O’Connell, S. Sinei, L. Maganga, H. Kibuuka, S. Nitayaphan, M. Rao, M. A Marovich, S. J. Krebs, M. Rolland, B. T Korber, G. M. Shaw, N. L Michael, M. L. Robb, S. Tovanabutra, J. H Kim, Rare HIV-1 transmitted/founder lineages identified by deep viral sequencing contribute to rapid shifts in dominant quasispecies during acute and early infection, PLoS Pathog. 13 (7) (2017) e1006510.

64. C. S. Oberle, C. Magnus, B. Joos, P. Rusert, D. Beauparlant, R. Kouyos, A. Trkola, K. J Metzner, H. F. Gunthard, Reply to correspondence ‘Conserved signatures indicate HIV-1 transmission is under strong selection and thus is not a “stochastic” process’ by Gonzalez et al., Retrovirology 2017, Retrovirology 14 (1) (2017) 14.

65. A. D. Redd, A. N. Collinson-Streng, N. Chatziandreou, C. E Mullis, O. Laeyendecker, C. Martens, S. Ricklefs, N. Kiwanuka, P. H Nyein, T. Lutalo, M. K Grabowski, X. Kong, J. Manucci, N. Sewankambo, M. J Wawer, R. H. Gray, S. F Porcella, A. S. Fauci, M. Sagar, D. Serwadda, T. C Quinn, Previously transmitted HIV-1 strains are preferentially selected during subsequent sexual transmissions, J. Infect. Dis. 206 (9) (2012) 1433–1442.

66. H. Schuitemaker, A. B. van’t Wout, P. Lusso, Clinical significance of HIV-1 coreceptor usage, J Transl Med 9 Suppl 1 (2011) S5.

67. M. J. Deymier, Z. Ende, A. E. Fenton-May, D. A Dilernia, W. Kilembe, S. A Allen, P. Borrow, E. Hunter, Heterosexual Transmission of Subtype C HIV-1 Selects Consensus-Like Variants without Increased Replicative Capacity or Interferon-I± Resistance, PLoS Pathog. 11 (9) (2015) e1005154.

68. J. E. Volk, J. L Marcus, T. Phengrasamy, D. Blechinger, D. P Nguyen, S. Follansbee, C. B Hare, No new hiv infections with increasing use of hiv preexposure prophylaxis in a clinical practice setting, Clinical Infectious Diseases 61 (10) (2015) 1601–1603. arXiv:/oup/backfile/content_public/journal/cid/61/10/10.1093/cid/civ778/2/civ778.pdf, doi: 10.1093/cid/civ778. URL http://dx.doi.org/10.1093/cid/civ778

69. E. Hoornenborg, M. Prins, R. C. A Achterbergh, L. R. Woittiez, M. Cornelissen, S. Jurriaans, N. A Kootstra, P. L. Anderson, P. Reiss, H. J. C. de Vries, J. M Prins, G. J. de Bree, Acquisition of wild-type HIV-1 infection in a patient on pre-exposure prophylaxis with high intracellular concentrations of tenofovir diphosphate: a case report, Lancet HIV.

70. C. Van Tienen, D. van de Vijver, T. Noori, A. Sonnerborg, C. Boucher, Letter to the editor: Pre-exposure prophylaxis for hiv in europe: The need for resistance surveillance, Eurosurveillance 22 (11).

71. D. R. Smith, N. Mideo, Modelling the evolution of hiv-1 virulence in response to imperfect therapy and prophylaxis, Evolutionary applications 10 (3) (2017) 297–309.

72. F. Zanini, J. Brodin, L. Thebo, C. Lanz, G. Bratt, J. Albert, R. A Neher, Population genomics of intrapatient HIV-1 evolution, Elife 4.

73. P. J. McLaren, C. Coulonges, I. Bartha, T. L Lenz, A. J. Deutsch, A. Bashirova, S. Buchbinder, M. N Carrington, A. Cossarizza, J. Dalmau, A. De Luca, J. J Goedert, D. Gurdasani, D. W Haas, J. T. Herbeck, E. O Johnson, G. D. Kirk, O. Lambotte, M. Luo, S. Mallal, D. van Manen, J. Martinez-Picado, L. Meyer, J. M Miro, J. I. Mullins, N. Obel, G. Poli, M. S Sandhu, H. Schuitemaker, P. R Shea, I. Theodorou, B. D Walker, A. C. Weintrob, C. A Winkler, S. M. Wolinsky, S. Raychaudhuri, D. B Goldstein, A. Telenti, P. I. de Bakker, J. F Zagury, J. Fellay, Polymorphisms of large effect explain the majority of the host genetic contribution to variation of HIV-1 virus load, Proc. Natl. Acad. Sci. U.S.A. 112 (47) (2015) 14658–14663.

74. T. D. Hollingsworth, O. Laeyendecker, G. Shirreff, C. A Donnelly, D. Serwadda, M. J Wawer, N. Kiwanuka, F. Nalugoda, A. Collinson-Streng, V. Ssempijja, W. P Hanage, T. C. Quinn, R. H Gray, C. Fraser, HIV-1 transmitting couples have similar viral load set-points in Rakai, Uganda, PLoS Pathog. 6 (5) (2010) e1000876.

75. F. Blanquart, C. Wymant, M. Cornelissen, A. Gall, M. Bakker, D. Bezemer, M. Hall, M. Hillebregt, S. H Ong, J. Albert, N. Bannert, J. Fellay, K. Fransen, A. J Gourlay, M. K. Grabowski, B. Gunsenheimer-Bartmeyer, H. F Gunthard, P. Kivela, R. Kouyos, O. Laeyendecker, K. Liitsola, L. Meyer, K. Porter, M. Ristola, A. van Sighem, G. Vanham, B. Berkhout, P. Kellam, P. Reiss, C. Fraser, Correction: Viral genetic variation accounts for a third of variability in HIV-1 set-point viral load in Europe, PLoS Biol. 15 (7) (2017) e1002608.

76. E. Hodcroft, J. D Hadfield, E. Fearnhill, A. Phillips, D. Dunn, S. O’Shea, D. Pillay, A. J. Leigh Brown, C. Aitken, D. Asboe, A. Pozniak, P. Cane, H. Castro, D. Dunn, E. Fearnhill, K. Porter, D. Chadwick, D. Churchill, D. Clark, S. Collins, V. Delpech, S. Douthwaite, A. M Geretti, A. Hale, S. Hue, S. Kaye, P. Kellam, L. Lazarus, A. Leigh-Brown, T. Mbisa, N. Mackie, C. Orkin, E. Nastouli, D. Pillay, A. Phillips, C. Sabin, E. Smit, K. Templeton, P. Tilston, D. Webster, I. Williams, H. Zhang, M. Zuckerman, J. Ainsworth, S. Allan, J. Anderson, A. Babiker, D. Chadwick, V. Delpech, D. Dunn, M. Fisher, B. Gazzard, R. Gilson, M. Gompels, P. Hay, T. Hill, M. Johnson, S. Kegg, C. Leen, F. Martin, M. Nelson, C. Orkin, A. Palfreeman, A. Phillips, D. Pillay, J. Pritchard, F. Post, C. Sabin, M. Sachikonye, A. Schwenk, A. Tariq, J. Walsh, The contribution of viral genotype to plasma viral set-point in HIV infection, PLoS Pathog. 10 (5) (2014) e1004112.

77. R. Sanjuan, M. R Nebot, J. B. Peris, J. Alcami, Immune activation promotes evolutionary conservation of T-cell epitopes in HIV-1, PLoS Biol. 11 (4) (2013) e1001523.

78. K. A. Lythgoe, C. Fraser, New insights into the evolutionary rate of HIV-1 at the within-host and epidemiological levels, Proc. Biol. Sci. 279 (1741) (2012) 3367–3375.

79. N. Pantazis, K. Porter, D. Costagliola, A. De Luca, J. Ghosn, M. Guiguet, A. M Johnson, A. D. Kelleher, C. Morrison, R. Thiebaut, L. Wittkop, G. Touloumi, Temporal trends in prognostic markers of HIV-1 virulence and transmissibility: an observational cohort study, Lancet HIV 1 (3) (2014) e119–126.

80. C. H. van Dorp, M. van Boven, R. J. de Boer, Immuno-epidemiological modeling of HIV-1 predicts high heritability of the set-point virus load, while selection for CTL escape dominates virulence evolution, PLoS Comput. Biol. 10 (12) (2014) e1003899.

81. H. M. Doekes, C. Fraser, K. A Lythgoe, Effect of the Latent Reservoir on the Evolution of HIV at the Within-and Between-Host Levels, PLoS Comput. Biol. 13 (1) (2017) e1005228.

82. B. Vrancken, G. Baele, A. M Vandamme, K. van Laethem, M. A Suchard, P. Lemey, Disentangling the impact of within-host evolution and transmission dynamics on the tempo of HIV-1 evolution, AIDS 29 (12) (2015) 1549–1556.

83. T. T. Immonen, T. Leitner, Reduced evolutionary rates in HIV-1 reveal extensive latency periods among replicating lineages, Retrovirology 11 (2014) 81.

84. I. Bartha, P. Simon, V. Muller, Has HIV evolved to induce immune pathogenesis?, Trends Immunol. 29 (7) (2008) 322–328.

85. E. Romero-Severson, H. Skar, I. Bulla, J. Albert, T. Leitner, Timing and order of transmission events is not directly reflected in a pathogen phylogeny, Mol. Biol. Evol. 31 (9) (2014) 2472–2482.

86. E. M. Volz, E. Romero-Severson, T. Leitner, Phylodynamic Inference across Epidemic Scales, Mol. Biol. Evol. 34 (5) (2017) 1276–1288.

87. F. Giardina, E. O. Romero-Severson, J. Albert, T. Britton, T. Leitner, Inference of Transmission Network Structure from HIV Phylogenetic Trees, PLoS Comput. Biol. 13 (1) (2017) e1005316.

88. J. L. Aldous, S. K Pond, A. Poon, S. Jain, H. Qin, J. S Kahn, M. Kitahata, B. Rodriguez, A. M Dennis, S. L. Boswell, R. Haubrich, D. M Smith, Characterizing hiv transmission networks across the united states, Clinical Infectious Diseases 55 (8) (2012) 1135–1143. arXiv:/oup/backfile/content_public/journal/cid/55/8/10.1093_cid_cis612/1/cis612.pdf, doi:10.1093/cid/cis612. URL +http://dx.doi.org/10.1093/cid/cis612

89. H. Heesterbeek, R. M Anderson, V. Andreasen, S. Bansal, D. De Angelis, C. Dye, K. T Eames, W. J. Edmunds, S. D Frost, S. Funk, T. D Hollingsworth, T. House, V. Isham, P. Klepac, J. Lessler, J. O. LloydSmith, C. J Metcalf, D. Mollison, L. Pellis, J. R Pulliam, M. G. Roberts, C. Viboud, N. Arinaminpathy, F. Ball, T. Bogich, J. Gog, B. Grenfell, A. L Lloyd, A. Mclean, P. O’Neill, C. Pearson, S. Riley, G. S Tomba, P. Trapman, J. Wood, Modeling infectious disease dynamics in the complex landscape of global health, Science 347 (6227) (2015) aaa4339.

90. P. Libin, E. Vanden Eynden, F. Incardona, A. Nowe, A. Bezenchek, A. Sonnerborg, A.-M. Vandamme, K. Theys, G. Baele, Phylogeotool: interactively exploring large phylogenies in an epidemiological context, Bioinformatics.

91. A. B. Abecasis, A. M Wensing, D. Paraskevis, J. Vercauteren, K. Theys, D. A. Van de Vijver, J. Albert, B. Asjo, C. Balotta, D. Beshkov, et al., Hiv-1 subtype distribution and its demographic determinants in newly diagnosed patients in europe suggest highly compartmentalized epidemics, Retrovirology 10 (1) (2013) 7.

92. P. Libin, G. Beheydt, K. Deforche, S. Imbrechts, F. Ferreira, K. Van Laethem, K. Theys, A. P Carvalho, J. Cavaco-Silva, G. Lapadula, et al., Regadb: community-driven data management and analysis for infectious diseases, Bioinformatics 29 (11) (2013) 1477–1480.

93. T. S. Group, et al., Global epidemiology of drug resistance after failure of who recommended first-line regimens for adult hiv-1 infection: a multicentre retrospective cohort study, The Lancet infectious diseases 16 (5) (2016) 565–575.

94. L. C. J Alcantara, S. Cassol, P. Libin, K. Deforche, O. G Pybus, M. Van Ranst, B. Galvao-Castro, A.-M. Vandamme, T. de Oliveira, A standardized framework for accurate, high-throughput genotyping of recombinant and non-recombinant viral sequences, Nucleic acids research 37 (suppl_2) (2009) W634–W642.

95. J. O. Wertheim, A. J. Leigh Brown, N. L Hepler, S. R. Mehta, D. D Richman, D. M. Smith, S. L. Kosakovsky Pond, The global transmission network of hiv-1, The Journal of infectious diseases 209 (2) (2013) 304–313.

